# SEDATION DIFFERENTIALLY AFFECTS DISTORTION-PRODUCT AND STIMULUS-FREQUENCY OTOACOUSTIC EMISSIONS IN CHINCHILLAS

**DOI:** 10.64898/2026.08.26.746474

**Authors:** Samantha Hauser, Andrew Sivaprakasam, Hari Bharadwaj, Michael Heinz

**Author notes:** Corresponding Author: Samantha Hauser.

## Abstract

**Purpose:** Otoacoustic emissions (OAEs) are used to assess outer hair cell (OHC) function. Clinical interpretation of OAE responses, however, is often limited to a present/absent binary since both physiological factors and measurement variability affect the measured OAE amplitude. Prior work showed elevated OAE responses in sedated compared to awake chinchillas, pointing to the potential influence of the medial olivocochlear (MOC) efferents on amplitudes, but this finding is inconsistent across species and OAE type. Here, we aimed to further investigate the effect of anesthesia on distortion-and reflection-type emissions in chinchillas using swept stimuli and more reliable calibration methods.

**Methods:** Swept distortion-product (DP) and stimulus-frequency (SF) OAEs were measured in chinchillas with and without ketamine/xylazine sedation. Stimuli were presented using in-ear forward pressure level calibrations. DPOAE and SFOAE amplitudes and estimated Q_erb_ from SFOAE group delays were compared across the two conditions.

**Results:** We found that low-frequency DPOAE amplitudes were elevated when animals were sedated. The difference in SFOAE amplitudes was more variable across animals but appeared mildly reduced in sedated animals. Q_erb_ estimates were slightly higher in sedated animals at some frequencies. The effect of sedation was not different across sexes.

**Conclusion:** Taken together, these findings suggest that sedation impacts OAE measurements in chinchillas. MOC modulation could account for the present findings and differences across species. For diagnostic precision, OAE responses should be considered in the context of not only intrinsic OHC function but also extrinsic physiological processes that can modulate OHCs.

## Introduction

Because cochlear and auditory health cannot be directly assessed in human patients, clinicians and researchers must rely on non-invasive physiological measures to evaluate auditory system function. Physiological measures that assess specific sites of lesion are especially critical for improving hearing diagnostics and for the development of targeted treatments for hearing loss. As site-of-lesion-specific treatments for hearing loss are developed (e.g. pharmaceuticals, gene therapy), identifying the appropriate candidates for intervention will be just as important as the effectiveness of the treatment itself. Animal models provide an important translational bridge for assessing the relationship between specific auditory lesions and physiological changes. However, species-specific effects and differences in testing paradigms between humans and animals limit direct translation and require cautious interpretation of results.

Otoacoustic emissions (OAEs) are a physiological measure commonly used to diagnose cochlear pathology. OAEs are sounds emitted from the cochlea that largely reflect outer hair cell (OHC) function [1, 2]. They are typically absent in individuals with more than a mild sensorineural hearing loss [3], and they are useful for hearing screenings in children, given that they can be collected passively and do not require a behavioral response. Many studies of cochlear pathologies in animal models have also relied on OAEs as an assay of OHC function. OAEs are absent in animal models with genetic mutations affecting OHCs (e.g., [4]) and are reduced or absent following noise exposure (e.g., [5]). In cases of noise-induced permanent threshold shifts, OAE amplitudes remain reduced, but they can recover alongside improvements in hearing thresholds for milder noise exposure paradigms [6, 7]. Exposures that leave OAE amplitudes unchanged are typically interpreted as having no effect on OHC function.

Despite their utility, many sources of variability have complicated the use of OAEs as a diagnostic for hearing loss. Although hearing thresholds are generally well correlated with OAE amplitudes given their shared sensitivity to OHC function, significant individual variability exists, especially in normal hearing ears [8]. Probe placement and standing waves in the ear canal introduce variability by altering sound level at the tympanic membrane. These differences in individual ear acoustics have been addressed with forward pressure level (FPL) and emitted pressure level (EPL) calibrations [8, 9]. These in-ear calibration techniques can reduce the variability of measurements, particularly at high frequencies. The interpretation of a given OAE amplitude is also dependent on the cochlear source of the emission, further complicating the relationship between OAEs and audiometric thresholds [10]. Distortion product otoacoustic emissions (DPOAEs) arise primarily from non-linear distortion in the cochlea, whereas stimulus-frequency otoacoustic emissions (SFOAEs) and transient-evoked otoacoustic emissions (TEOAEs) are generated by linear coherent reflection [11]. Recent work has shown that these emission types are differentially affected by specific cochlear pathologies [12]. Because different cochlear phenomena drive these emissions, not all changes in OHC function necessarily result in a uniform effect across otoacoustic emission types. These differences across OAE types appear to provide useful insight into the underlying cochlear pathologies when used together [13].

Additionally, OHCs receive efferent input from the medial olivocochlear system (MOC), which modulates OAE amplitudes by altering the gain of OHCs [14]. Contralateral acoustic stimulation has been shown to reduce both DPOAE and SFOAE amplitudes in some individuals (e.g., [15]). The status of the middle ear as well as the strength of the middle-ear muscle reflex (MEMR) can also contribute to OAE amplitudes measured in the ear canal. Activation of the MEMR changes the immittance properties of the middle ear, which could alter the level of the OAE stimulus reaching the cochlea as well as alter the level of the back-propagated emission, especially in the low frequencies.

These physiological sources of variability are often ignored in the interpretation of OAE amplitudes. A reduction in OAEs could correspond to an increase in MOC or MEMR strength, and a decrease in OHC function could be hidden by weakened efferent feedback. In fact, change in drive to the efferent pathways is a likely source of the finding that chinchillas with carboplatin-induced inner hair cell (IHC) dysfunction show subtle but consistent increased OAE amplitudes [16–18].

Prior studies have also investigated the role of the efferent system by examining OAEs under sedation, which alters efferent feedback. However, results across species are variable. While studies in chinchillas have shown that TEOAEs and DPOAEs are higher during sedation than when the same animals were awake [19], most studies show either no effect of sedation on human OAE amplitudes [20, 21] or an overall decrease in OAE amplitudes [22, 23]. Calibration differences and variation from probe placements diminish the precision of these measurements and have made identifying these subtle effects difficult to confirm. Forward pressure level calibrations that account for these differences are likely to allow for measurements more sensitive to the physiological modulation of OHC function and the corresponding change in OAE amplitudes.

In this study, we expand upon these prior studies of the effect of sedation on OAEs in chinchillas, taking advantage of improved calibration techniques, more comprehensive OAE paradigms with swept-frequency stimuli, and both distortion and reflection OAE types. To ensure that subtle effects could be detected and to minimize variability from the probe placement and ear canal acoustics [8, 9], we used forward pressure level calibration in conjunction with frequency-sweeping stimuli to allow for dense sampling of the OAEs [24, 25]. We also utilized both SFOAE and DPOAEs to assess both mechanisms of OAE generation, linear coherent reflection and nonlinear distortion [11]. Ultimately, this work contributes to our understanding of the diagnostic precision of OAEs.

## Methods and Materials

### Subjects

Twenty young chinchillas (*Chinchilla lanigera*), 6 to 36 months of age (mean 13.6, SD 9.5; male: n = 9; female: n = 11) were used in this study. The animals were housed with a 12/12 h light/dark cycle in a temperature and humidity-controlled facility. They were provided daily, non-auditory enrichment. All procedures were approved by the Purdue University Animal Care and Use Committee (Protocol #1110000123).

All chinchillas were confirmed to have normal hearing via sedated auditory brainstem response (ABR) testing at 0.5, 1, 2, 4, and 8 kHz [26]. Thresholds were determined via a cross-correlation method based on ABRpresto [27, 28]. All data collection was completed in a double-walled sound booth. OAEs and ABR threshold estimates were only measured in the right ear.

### Awake Restraint

Data collection occurred in two experimental conditions: awake restraint or under sedation. Awake measurements were obtained while the animals were alert and positioned comfortably in a custom restraint device (Online Resource 1) used in previous studies [6] and modified based on [29]. Animals were acclimated to the restraint tube prior to testing. Most awake test sessions were 30-60 minutes but were terminated if the animal failed to participate or did not tolerate the eartip, in which case testing resumed after a break or on another day. Awake and sedated sessions were completed within two weeks of each other.

### Anesthesia

Animals were sedated with xylazine (4 mg/kg, s.c.) followed by ketamine (20-40 mg/kg, s.c.) approximately 10 minutes later. The animal’s temperature was monitored throughout the procedure with a rectal probe and maintained at 37°C using a feedback-controlled heating pad (Harvard Apparatus, 50-7053F). Heart rate and SpO_2_ were also monitored during the session with a pulse oximeter (Nonin 8600V, Plymouth, MN) on one of the paws. Eye ointment was placed to prevent dryness and corneal injury, and the head was elevated using a stereotaxic bite bar to optimize airway patency. Supplemental oxygen was provided throughout recordings with an oxygen tube placed near the animal’s nose. After data collection was completed, atipamezole (0.4 mg/kg, i.p.) was administered to facilitate recovery. Animals were monitored closely for at least three days after anesthesia to ensure good recovery. 6 mL of lactated Ringer’s solution was administered both prior to induction and after completion of recording to maintain hydration. Chinchillas were sedated for 2–4-hour sessions, which included other electrophysiological testing (ABR threshold, envelope following responses). Sedated OAE measures were typically taken toward the end of an experimental session.

### Equipment and Calibration

All stimuli were presented through electrically shielded ER2 insert earphone transducers with a foam or rubber tip coupled to an ER-10B+ microphone system (Etymotic). At the beginning of each day of data collection, a probe calibration to estimate the Thevenin-equivalent source pressure and impedance needed for FPL calibrations was completed. The ER-10B+ probe was coupled to custom brass cylinders of five different lengths, each with an 8 mm diameter [30] (Online Resource 2). By measuring the acoustic response to clicks at the microphone in each of the tubes, where acoustic impedance values can be approximated, estimates of source pressure and impedance of the probe were determined by minimizing the “calibration error”. This unitless metric is an energy ratio averaged over 2-8 kHz and scaled by 10000. This “error” was calculated for each channel and was always below 10 prior to running an in-ear calibration [6, 31]. Prior to data collection in each animal, an in-ear calibration using the same click stimulus was conducted to measure the immittance properties of the individual ear for each probe placement. Coefficients for a 256-tap inverse FIR filter were then estimated from the response of the ear to create a flat FPL response at the tympanic membrane. The coefficients were applied to custom Tucker-Davis Technologies (TDT) hardware so that an attenuation of 0 dB would produce a flat 105 dB stimulus up to 10 kHz. Programmable attenuators (PA5, TDT) were then used to adjust the sound level to the desired output. All stimuli were presented to the right ear only. Data were acquired with a 48.828 kHz sampling rate.

### Distortion Product Otoacoustic Emissions (DPOAEs)

DPOAEs were presented with a frequency ratio of F_2_/F_1_ = 1.22 and F_1_ and F_2_ levels of 75 and 65 dB FPL, respectively. F_2_ was swept upward from 0.5-16 kHz at a rate of 1 octave per second. A signal-to-noise ratio (SNR)-based data collection method adapted from [12] was used to make data collection more efficient. During data collection, the OAE amplitude and noise levels were calculated at half-octave bands centered at 707 to 8000 Hz. Individual trials with amplitudes exceeding three standard deviations above the median were excluded from the minimum trial count. Data collection continued until at least 12 accepted trials were obtained and an SNR of at least 6 dB was reached at each of the specified frequencies. A 50-trial limit was implemented to minimize animal discomfort in cases where the SNR or minimum number of clean trials criteria were not met. All raw trials were saved independently, even those that did not meet the initial acceptance criteria.

DPOAEs were analyzed offline using custom MATLAB scripts based on the recorded microphone response of each trial. First, individual trials were screened for artifacts as in the data collection procedure. A least-squares fit (LSF) was applied to each individual trial to estimate the magnitude of the distortion product (DP; 2F_1_-F_2_) at each of 512 logarithmically spaced frequency points from 0.5 to 16 kHz. For each F_2_ frequency point, the microphone response within a 250 ms Hann-tapered window centered on the time in which that frequency occurred in the sweep was fitted to a model containing the sine and cosine components of the expected DPOAE response (2F_1_-F_2_) akin to [24, 25, 32]. The fitted coefficients of the components define the complex pressure of the DP, from which the magnitude and phase of the DPOAE level was calculated. Noise floor estimates were made using the same least-squares fitting procedure, but with the response fit to four nearby frequencies (0.90, 0.88, 0.86, 0.84 times the DP frequency). Prior to analyzing the average response across trials, artifacts were removed by performing the LSF analysis on each trial individually. The median and standard deviation across trials was computed at each frequency. Any trial-level estimate of the OAE amplitude that exceeded the median by more than three standard deviations was considered noise. These “noisy” points were then removed from the original trial in the time-domain by eliminating a 300 ms window (250 ms window ± 25 ms) centered around the time when that frequency occurred in the stimulus. The screened trials were then averaged to generate a single, clean time-domain response which was used to obtain the final DP amplitude and noise floor with the LSF procedure. DPOAE amplitudes were converted to emitted pressure level (EPL) using the in-ear calibration data. To summarize the frequency response across the continuous sweep, an SNR-weighted average was calculated at nine half-octave-spaced points with center frequencies from 707 to 11300 Hz.

### Stimulus-Frequency Otoacoustic Emissions (SFOAEs)

SFOAEs were measured using a suppression paradigm [24]. Each trial consisted of three stimulus conditions: probe alone (P), suppressor alone (S), and probe plus suppressor (PS). The probe stimulus level was 40 dB FPL and was swept downward in frequency from 16 to 0.5 kHz. The suppressor tone was presented 20 dB higher in level than the probe and swept 50 Hz lower in frequency. Both the probe and the suppressor were swept at a rate of 1 octave per second. To minimize stimulus artifact contamination, the phase of the suppressor was rotated by 180 degrees across successive trials. The SFOAE response (i.e., the residual) was calculated by subtracting the combined probe plus suppressor response from the sum of the individual probe and suppressor responses (i.e., SFOAE = P + S − PS).

The initial stages of the SFOAE analysis were similar to DPOAE analysis, but the analysis was performed on the SFOAE residual (P+S-PS) of each trial rather than the raw microphone response. As with DPOAEs, noisy windows during individual trials were excluded from the average response. The LSF procedure was used to calculate the complex response of the SFOAE and the noise. The analysis window varied continuously from 0.16 seconds at 0.5 kHz to 0.038 seconds at 16 kHz to keep the number of phase rotations in each analysis window similar [12, 33]. The frequency-domain SFOAE was transformed to the time domain by inverse Fourier transform, rectangularly windowed from 0 to 21 ms, and transformed back; this procedure aids in reducing noise and long latency reflections. The noise was transformed with this same process, with the response fit to models of 1.1, 1.12, 1.14, 1.16 times the test frequency. The resulting swept SFOAE response is peaky, and previous work shows the points at local maxima provide better estimates of phase-gradient than averaging across the entire response [34]. To this end, amplitude and phase calculations only used the peaks of the SFOAE as well as the two points adjacent to a peak (one on either side). The amplitudes and Q_erb_ were summarized at nine half-octave bands with center frequencies between 707 and 11300 Hz. A simple average of the points from the peaks was calculated for each band. Q_erb_ was calculated in each band by taking the slope of the unwrapped phase versus frequency function to get the phase-gradient delay, which was converted to periods (N_sfoae_) by multiplying by frequency. N_sfoae_ was then multiplied by a dimensionless, species-invariant conversion factor, r, (r = 1.25 as derived by [35]) to estimate Q_erb_.

### Statistical Analyses

DPOAE amplitudes, SFOAE amplitudes, and SFOAE Q_erb_ estimates were analyzed using linear mixed effects models using lme4 [36], in R. Each outcome was modeled as a function of frequency (nine half-octave bands treated as a categorical fixed effect), sedation state (awake vs sedated), and their interaction, with subject as a random effect. Overall effects were assessed with Type II Wald F tests with Kenward-Roger degrees of freedom (car package [37]). Awake vs sedated contrasts were estimated at each frequency using estimated marginal means (emmeans package [38]). Per-frequency contrasts are reported without correction for multiple comparisons. p < 0.05 was used as the threshold for statistical significance and tests are two-tailed.

## Results

### Emissions from an individual animal

All animals used in this study showed robust otoacoustic emissions across the frequency range tested (0.5-16 kHz) for both OAE types. DPOAEs were consistently well above the noise floor in all animals with an average amplitude of approximately 35 dB EPL across frequencies and conditions. DPOAE amplitudes were generally highest in the mid to high frequencies (4-10 kHz). SFOAEs were overall lower in amplitude than DPOAEs with an average amplitude across conditions and frequencies of 13 dB EPL. In chinchillas, DPOAEs were consistently larger in amplitude than SFOAEs, consistent with responses in rodents, but opposite of findings in humans [39]. SFOAEs also exhibited more pronounced fine structure than DPOAEs, with regular dips in the response amplitude.

Figure 1 shows the DPOAE (a) and SFOAE (b) responses from a representative subject at 512 analysis points across the frequency range in the awake (red) and sedated (blue, dashed) conditions. For this subject, DPOAEs are clearly elevated under sedation compared to when the animal is awake. The effect is most prominent in the low frequencies, but DPOAEs are also slightly higher between 6-10 kHz. The effect of sedation is less clear in this animal’s SFOAEs; amplitudes are similar or slightly lower at many frequency points above 1.6 kHz in the sedation condition, but equal or greater below 1.6 kHz.

**Fig. 1.**
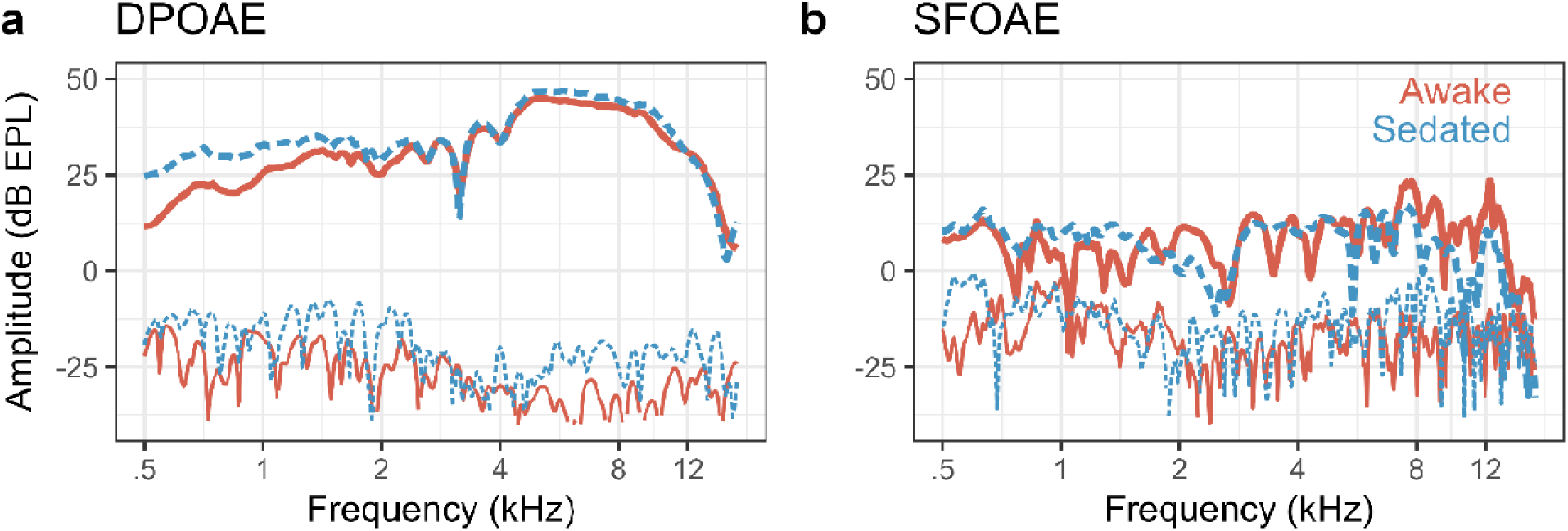
DPOAE (a) and SFOAE (b) amplitudes measured while a representative naïve, male chinchilla was awak (solid red) or sedated (dashed blue). The noise floor for each measure is shown with thinner lines. Fine structure of the response is largely repeatable across conditions. DPOAE amplitudes are higher when the animal is sedated than when it is awake; SFOAE amplitudes are higher at approximately 1.6 kHz and below but lower at frequencies above that in the sedated condition.

Other than the change in amplitude, features of both OAE types are largely stable between the two conditions. This subject shows a strong dip in the DPOAE response around 3 kHz which is consistent across both awake and sedated measurements. This repeatability of fine structure was seen in most animals tested.

### Low frequency DPOAE amplitudes are elevated following sedation

Figure 2a shows the average awake and sedated responses for 20 subjects at the nine half-octave bands. Figure 2b shows the within-animal difference between awake and sedated responses. A linear mixed effects model revealed that DPOAE amplitudes were significantly affected by frequency (F_8,323_ = 69.7, p < 0.0001) and sedation (F_1,323_ = 40.9, p < 0.0001), with a significant interaction between these two predictors (F_8,323_ = 3.12, p = 0.002). Post-hoc comparisons of awake and sedated conditions were conducted for each frequency band. The average sedated DPOAE amplitude was higher than the awake amplitude at every frequency point but only reached statistical significance in the lowest four frequency bins (0.7-2 kHz).

**Fig. 2.**
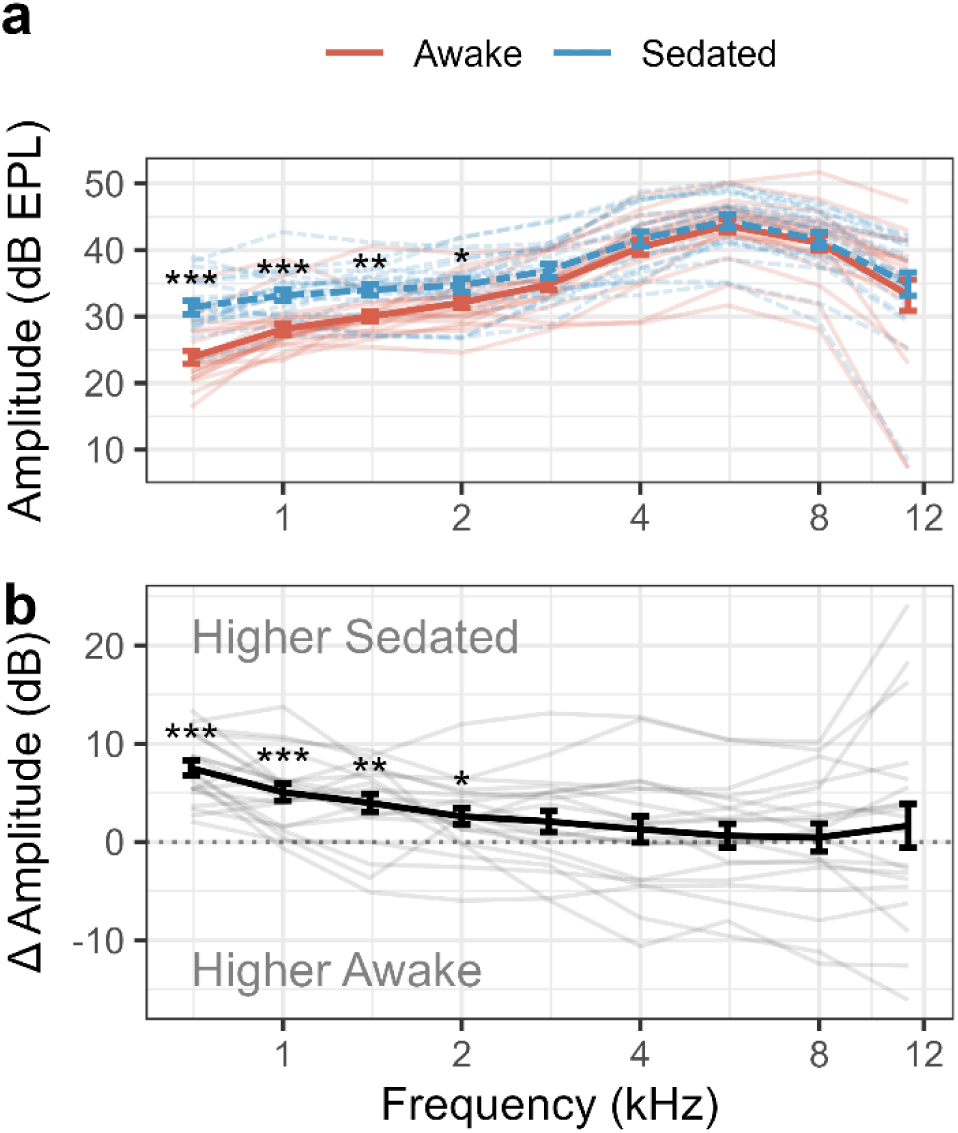
DPOAE amplitudes are higher in the low frequencies when animals are sedated compared to when they are awake. (a) DPOAE responses in animals while either awake (red) or sedated (blue, dashed). (b) The difference i amplitudes (Sedated - Awake) for individual animals. Thicker, bold lines show the average response and thin, light lines show individual animals. Error bars show S.E.M. * p < 0.05, ** p < 0.01, *** p < 0.001

As seen in Figure 2b, although every animal showed higher sedated DPOAEs than awake DPOAEs at 707 Hz, the change in amplitude was variable across animals. Some chinchillas demonstrated sedation-related enhancement only in the low frequencies, while others exhibited increased amplitudes across the entire frequency range.

We also tested a mixed effects model which additionally included a random slope for sedation by subject, thus allowing the effect of sedation to differ across subjects rather than assuming each showed the same sedation effect. In this model, the average effect of sedation is tested against between-subject variability. This is a stricter test that reduces the degrees of freedom and evaluates the effect across animals rather than all individual measurements. In this model, frequency (F_8,304_ = 95.3, p < 0.0001), sedation state (F_1,19_ = 7.73, p < 0.01), and their interaction (F_8,304_ = 4.26, p < 0.0001) remain significant. Post-hoc contrasts at individual half-octave bands yielded comparable results to the simpler model with larger DP amplitudes at low frequencies (0.7-1.4 kHz).

### SFOAE amplitudes are unchanged or reduced with sedation

As with DPOAEs, an average SFOAE amplitude was calculated for each half-octave window. However, SFOAEs were lower in amplitude than DPOAEs and some frequency points were near the noise floor. Because only the responses at peaks with at least a 6 dB SNR were counted toward the final response, some animals have no response in certain frequency windows. Those data were eliminated from further analysis (one data point from each of four animals: two points at 0.7 kHz and one point at each 1 and 1.4 kHz).

Figure 3 shows the average and individual SFOAE amplitudes across frequencies and sedation conditions. At the group level, sedation had a smaller and opposite effect on SFOAE amplitudes than on DPOAEs. SFOAE amplitudes were significantly affected by frequency (F_8,319_ = 15.3, p < 0.0001) and sedation (F_1,319_ = 5.84, p = 0.016). There was not a significant interaction between frequency and sedation (F_8,319_ = 0.99, p = 0.44). The average difference between awake and sedated SFOAEs was less than 3 dB at most frequencies. Comparisons at individual frequencies showed awake emissions were higher at 8 kHz (4.1 dB, t(319) = 2.04, p = 0.045), and at 11.3 kHz (5.3 dB, t(319) = 2.58, p = 0.010). Averaged SFOAE amplitudes were also 2.7 dB higher at 2 kHz and 1.6 dB higher at 5.6 kHz in the awake condition, but these effects were not significant.

**Fig. 3.**
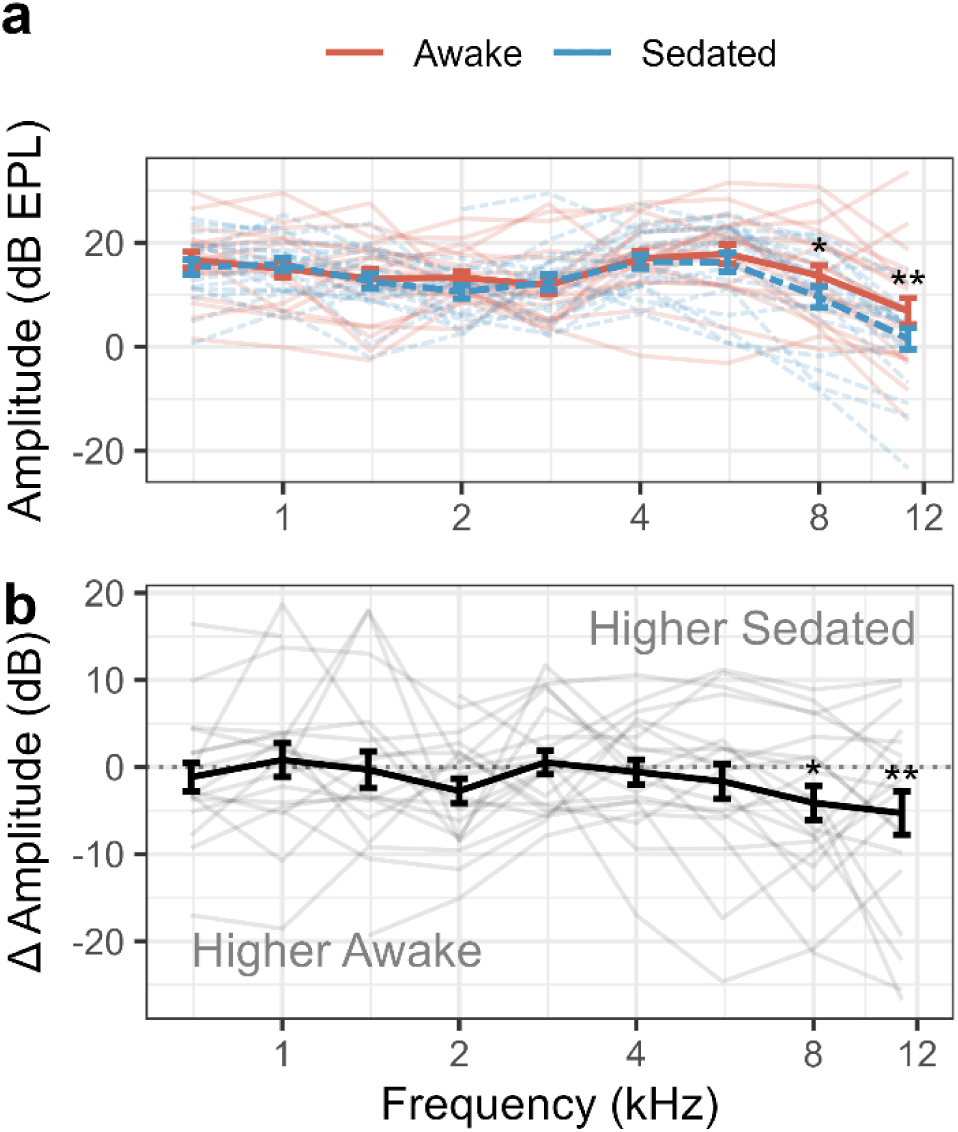
SFOAE amplitudes are unchanged at 5.6 kHz and below but lower with sedation at 8 and 11.3 kHz. (a) SFOAE responses in animals while either awake (red) or sedated (blue, dashed). (b) The difference in amplitudes (Sedated - Awake) for individual animals. Colors, error bars, and significance stars are as in Fig. 2.

Overall, there is more individual variability in the effect of sedation for the SFOAEs than for DPOAEs. Although there is a general trend to see a reduction in emission amplitudes in the high frequencies (Figure 3b), this is not true for every animal. Some showed higher sedated SFOAEs across all frequencies, some showed reductions, and many had variable effects across frequency.

### Cochlear tuning (Q_erb_) estimated from SFOAEs is sharper with sedation

In addition to SFOAE amplitudes, we also explored whether SFOAE-based estimates of cochlear tuning were affected by sedation. Tuning of a cochlear filter can be described by the equivalent rectangular bandwidth (ERB). Frequency selectivity, or the sharpness of a cochlear filter, at a given characteristic frequency (CF) can be represented by Q_erb_, where Q_erb_ is equal to CF divided by the ERB at that CF; sharper filters result in a higher Q_erb_. Auditory filters can be measured directly from auditory nerve fiber response [40], behaviorally [41], or from the phase of the SFOAE [35]. Here, Q_erb_ was calculated from the peaks of the response that were used to calculate SFOAE amplitude within each frequency band [34]. Q_erb_ as a function of frequency is plotted in Figure 4. In both conditions, Q_erb_ increases with increasing frequency from about 1.5 at 707 Hz to 10 at 11.3 kHz, as expected in normal hearing animals [35, 40, 42]. The average sedated cochlear-tuning estimates in Figure 4a (blue) are higher (i.e. sharper) at some frequencies than the average awake response (red). A linear mixed effects model predicting Q_erb_ reveals a significant effect of frequency (F_8,319_ = 47.2, p < .0001) and sedation (F_1,319_ = 7.3, p = 0.007). The interaction between sedation and frequency (F_8,319_ = 1.27, p = 0.26) was not significant. Although the interaction was not significant, the sedation effect appears more often at higher frequencies with Q_erb_ estimates significantly higher at 8 kHz (t(319) = 2.6, p = 0.010) and 11.3 kHz (t(319) = 2.0, p = 0.045). These changes, a 2.3 and 1.8 increase in Q_erb_ in the sedated condition at 8 and 11.3 kHz, indicate an approximately 25% and 16% decrease in ERB, respectively. Similarly, Q_erb_ with sedation is higher by 1.6 and 1.4 in sedated animals at 2 kHz (t(319) = 1.9, p = 0.059, a 44% decrease in ERB) and 4 kHz (t(319) = 1.7, p = 0.099, a 22% decrease in ERB) but does not reach statistical significance. The difference at each of the remaining frequencies was less than 0.3.

**Fig. 4.**
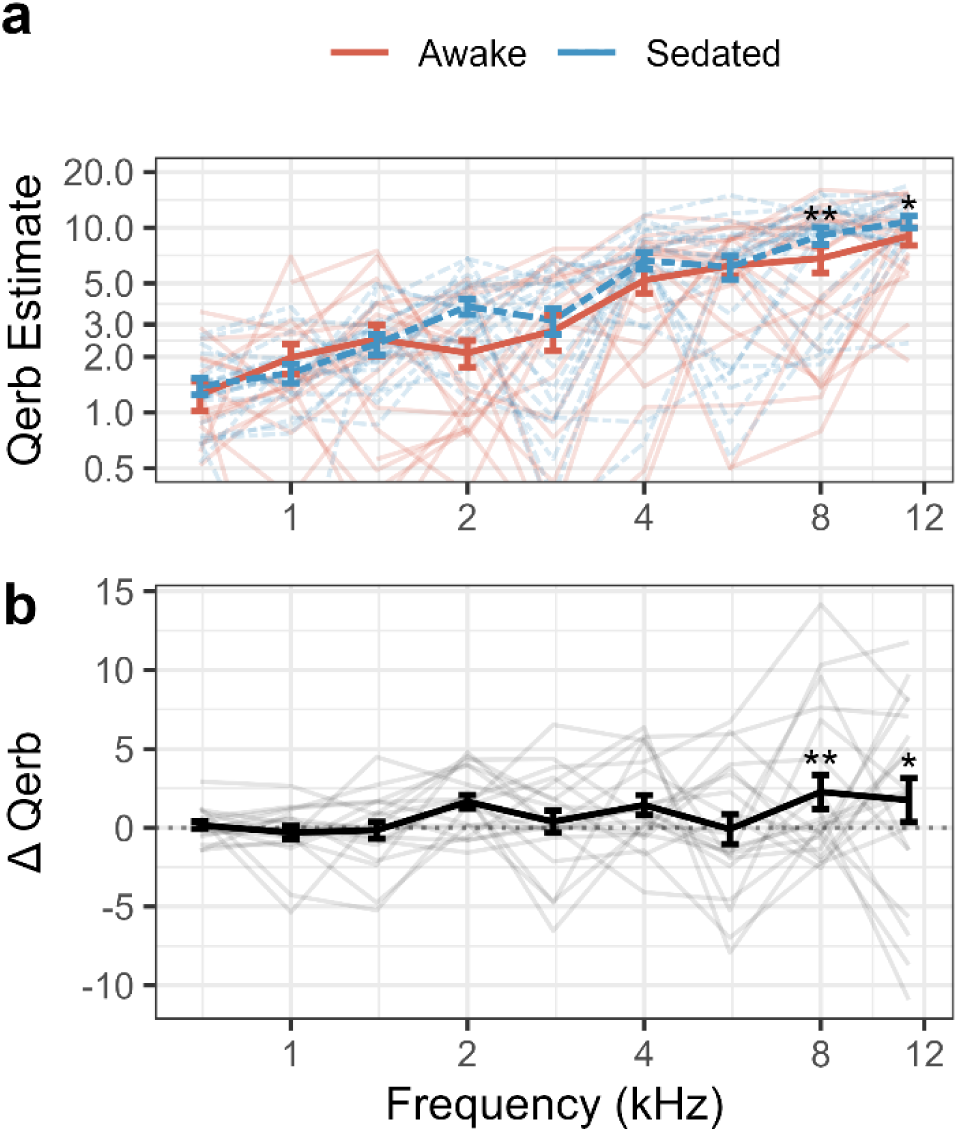
Cochlear tuning sharpness (Q_erb_) estimated from the phase of the SFOAE is increased in the high frequencies in sedated animals compared to awake. (a) Awake (red) and sedated (blue, dashed) results are shown as a function of center frequency. (b) The difference between sedated and awake Q_erb_ estimates within individual animals. Points above the dashed line represent sharper tuning when the animal was sedated compared to when the animal was awake. Colors, error bars, and significance stars as in Fig. 2.

### Effect of sedation on DPOAEs and SFOAEs does not differ across sexes

Sex is a potential source of variability in the responses across animals. The difference plots of DPOAE amplitudes (Figure 2), SFOAE amplitudes (Figure 3) and Q_erb_ (Figure 4) are replotted in Figure 5 with male and female subjects in different colors. The trends for each metric are consistent with the group data, though there are slight differences in responses between males and females. Females show a slightly greater increase in low frequency DPOAE amplitudes with sedation than males do (Figure 5a), but somewhat less of an effect than males in SFOAE amplitudes (Figure 5b) and Q_erb_ estimates (Figure 5c). Incorporating sex into the models predicting OAE amplitudes or Q_erb_, however, revealed no significant effect of sex for any measure (DP: F_1,18_ = 0.91, p = 0.35; SF: F_1,18_ = 1.7, p = 0.21; Q_erb_: F_1,18_ = 1.9, p = 0.19).

**Fig. 5.**
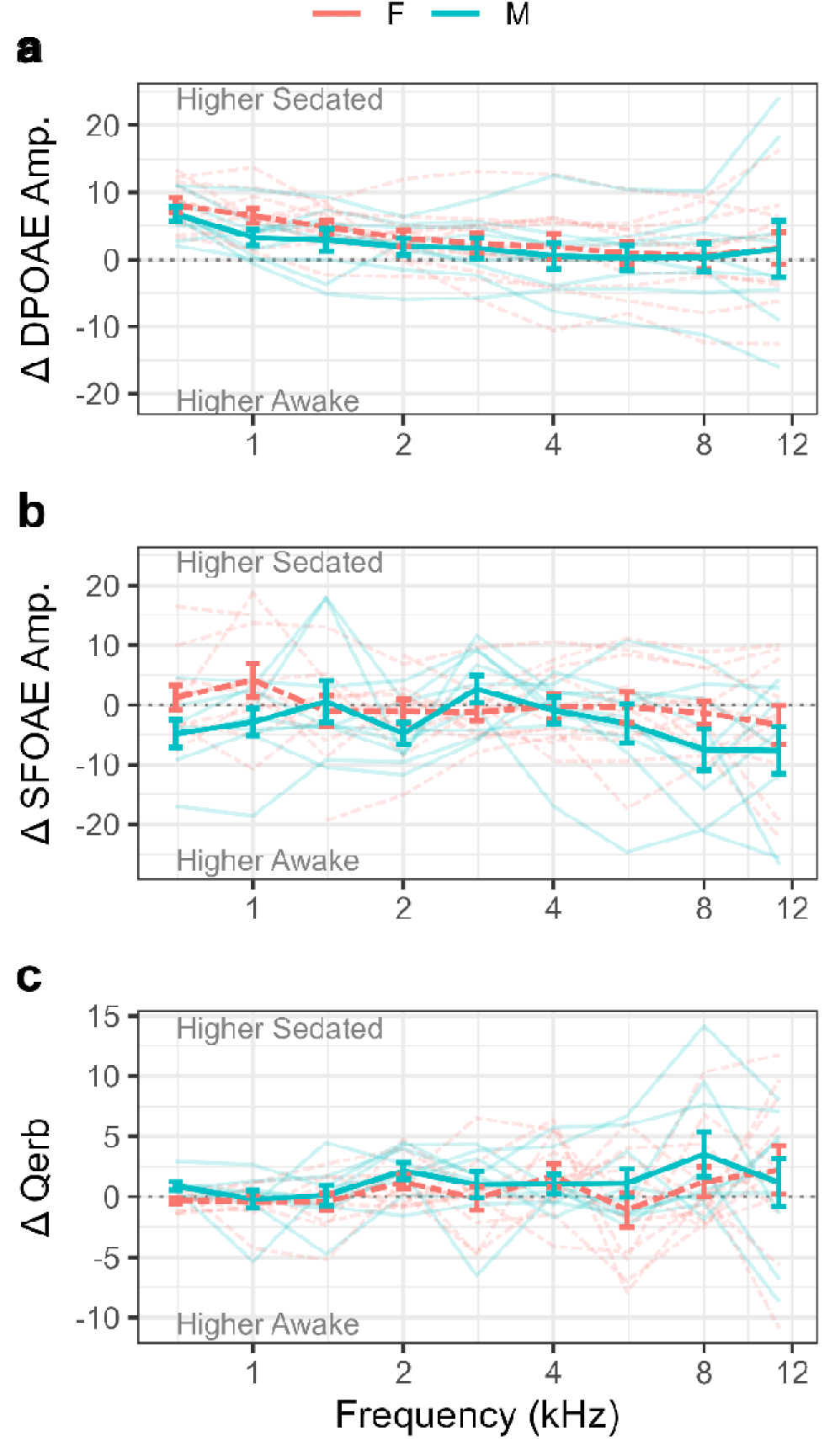
No statistically significant differences were observed between sexes. The effects of sedation on (a) DPOAE amplitudes, (b) SFOAE amplitudes, and (c) Q_erb_ estimates are not different across male and female animals. Data from males (n = 9) are in teal, data from females (n = 11) are in pink.

## Discussion

This study evaluated the effect of sedation on DPOAE and SFOAE amplitudes as well as cochlear tuning estimates (Q_erb_) from SFOAE phase in chinchillas. Overall, we find that our OAE measurements are qualitatively consistent with previous findings in sedated chinchillas [35], though prior reports generally did not compare awake and sedated conditions directly. The average Q_erb_ estimates are also consistent with previous findings in chinchillas [35]. We found that DPOAEs are elevated in the low frequencies when chinchillas are sedated with ketamine/xylazine relative to when they are awake. SFOAE amplitudes were lower in the high frequencies when animals were sedated, and Q_erb_ estimates were higher (i.e., suggesting sharper tuning) at some frequencies during sedation. Sex was not a significant predictor of OAE amplitudes or tuning in our data set, though subtle differences may be more readily identified in a larger sample of animals. Though the effect of time within the sedation window was not explored systematically in this study, data from one animal (Online Resource 3) suggests that differences in OAE amplitudes and estimates of cochlear tuning throughout the course of a two-hour sedated procedure are minimal.

The present report is most similar to the study by Harel and colleagues [19], which evaluated DPOAEs and TEOAEs in chinchillas before and after sedation with ketamine or sodium pentobarbital. Although a different DPOAE stimulus was used (L_1_=L_2_=50 dB SPL), they found that ketamine sedation resulted in significantly elevated DPOAEs at 1–1.7 kHz, 2.7 kHz, and 4–5.3 kHz. TEOAEs were also elevated at 1 kHz compared to pre-sedation. Sodium pentobarbital had no effect on DPOAEs and a slight increase in the TEOAEs at 1kHz. They also included several control experiments including sectioning of the middle ear muscles, saline injections, and repeated measures in the same condition, which showed no systematic effects.

The effect of anesthesia on DPOAE and TEOAEs has been studied in a number of species and across different anesthetic drugs. In humans, anesthesia generally results in a decrease in emission amplitudes [20, 21, 23], though the precise effect depends on the emission type (DPOAE or TEOAE) and the anesthesia used (e.g., propofol, isoflurane, sevoflurane, nitrous oxide, midazolam). Studies in rats and mice also tend to show a decrease in amplitudes. Isoflurane, and to a greater extent ketamine, resulted in an increase in DPOAE thresholds in C57BL/6J and C129/SvEv mice [43]. [44] found no effect of ketamine/xylazine on DPOAEs in rats but saw a reduction in emissions with isoflurane. A study of Sprague-Dawley rats found ketamine, especially in higher doses, reduces DPOAE and TEOAE amplitudes, attributing this effect to increased negative pressure in the middle ears during sedation [45]. In chickens and starlings, halothane and other anesthetics completely abolished DPOAEs [46]. In contrast, bats, like chinchillas, seem to consistently show an increase in OAE amplitudes with sedation. In the mustached bat, ketamine led to an increase in DPOAEs with greatest change at very high frequencies (>40 kHz) as well as a change in the optimal stimulus ratio [47].

The effect of sedation on OAEs in chinchillas is consistent with our hypothesis that the activity of the MOC alters DPOAE amplitudes and cochlear tuning. SFOAE amplitudes are also affected by the status of the OHCs [48], but the effect seen here is small and when present appears to be in the opposite direction of DPOAEs (lower amplitudes while sedated). The method of measuring SFOAEs could explain this. The SFOAE amplitudes presented here are not a direct measure of the OAE itself but of the SFOAE residual calculated by subtracting the response of the combined probe and suppressor trial (PS) from the sum of the individual stimulus (P + S) trials. Thus, if the change in cochlear physiology caused by sedation results in a change in the effectiveness of the suppressor, more (or less) of the probe-evoked OAE may remain in the PS trials and be subtracted from the true OAE when calculating the residual OAE, artificially reducing the amplitude. Since tuning narrows slightly with sedation, the probe-stimulus OAE is likely less suppressed than when the animal is awake. Thus, it appears likely that the optimal stimulus parameters for the measurement of SFOAEs using the suppressor paradigm may differ depending on whether the measurement is done awake or anesthetized. SFOAE phase measurements, as well as the derived tuning estimates, may be more informative than SFOAE amplitudes.

The species-specific effect of anesthesia on OAEs also supports the MOC hypothesis. Chinchillas are exceptional among most mammals in that the majority (∼80%) of their MOC fibers project contralaterally [49]. Since the MOC reflex arc crosses the midline of the brainstem twice, these fibers innervate the cochlea that was stimulated, so this strong contralateral bias predicts a larger effect for ipsilateral rather than contralateral stimulation. Chinchilla MOCs also show a bias toward the apex, consistent with our finding of the greatest effect of sedation on DPOAEs at low frequencies [49]. Thus, if the OAE stimulus activated the MOC pathway in the awake condition, the apparent elevation in OAEs with sedation may not only be due to an “unmasking” of the response, but a further depression of the emission when recorded in awake chinchillas. Mustached bats, which show similar increases in OAE amplitudes with sedation as chinchillas [47], also show this apical bias of MOC fibers and the higher distribution of crossed MOC fibers indicating that the specific neuroanatomical structure of the efferent system contributes to the effect of sedation on OAEs.

Though the findings of this study and others are consistent with MOC effects, we cannot rule out the contribution of the MEMR. Given the high sound level of the DPOAEs (75/65 dB FPL), it is possible that the awake measurements not only include the effect of the MOC reflex but also of the MEMR. The probe stimulus is softer for SFOAEs than DPOAEs, so there may be less contamination from the MEMR in this response. The frequency-specific effects, however, may provide some insight into the more likely mechanism of the observed effects. The finding of increased low-frequency DPOAEs under sedation is only partially consistent with the effect of the MEMR. The contraction of the MEMR while the animal is awake could reduce the low-frequency admittance of the middle ear system, an effect that would be absent while the animal is sedated. However, the shift in resonance to higher frequencies with contraction of the middle ear muscle would produce an alternating pattern of increases and decreases beyond the first resonance, which was not seen in our data. The increase in high frequency Q_erb_ is also more aligned with MOC-related changes in high-frequency gain than MEMR effects. Prior animal studies specifically investigating the MOC system as a potential mechanism of this effect have generally bisected the tensor tympani and stapedius muscles to eliminate the MEMR entirely [50]. This was not done in this study.

Placement of the probe may also vary to some extent in awake and sedated animals. Deep insertion of the eartip is easier in sedated animals and the position can be better maintained since the animal does not move during testing. Although awake chinchillas are seated in a restraint device designed to limit movement during awake testing, they are not fully immobilized. Their movements during or after probe placement while awake may lead to systematic differences between conditions; however, this difference is unlikely to be sufficient to explain these results. The FPL/EPL calibration technique employed here is designed to compensate for these differences in probe placement, unlike standard calibrations. Furthermore, the compressive growth function of DPOAEs suggests that at high levels, a change in stimulus level would likely lead to a smaller change in OAE amplitude. We would also expect that a reduction in stimulus level would have affected both DPOAEs and SFOAEs, but the observed differences were primarily related to DPOAEs.

### Implications for Precision Diagnostics

These results demonstrate that our diagnostic tests reflect not only the function of one cochlear component but the broader auditory system in which it is embedded. OAEs are an index of outer hair cell function, but they are modified by the efferent system, middle ear immittance, and physiological state of the subject, including sedation. For mechanistic, within-subject work, when a single species is measured in the same condition, these confounding effects are less likely to impact our interpretation that OHC status is the primary driver of OAE amplitudes. In human populations where testing is primarily done awake, the effect of the MOC may be present and variable across individuals, but may or may not impact findings. However, consideration of the diagnostic precision of our metrics becomes more impactful as the field moves towards more detailed measurement of auditory function and individualized characterization of cochlear dysfunctions. The additional confounds of sedation and differences in neuroanatomy must be considered in translating results from animal models to human subjects. Thus, measurement of these subtle effects and accounting for the many intrinsic and extrinsic factors that influence our diagnostic tests are critical to advancing precision audiology.

## Declarations

## Acknowledgements

This work was supported by the National Institutes of Health National Institute on Deafness and Other Communication Disorders grants F32DC021345 (SH), F30DC020916 (AS), T32DC016853 (SH, AS), R01DC022670 (HB), and R01DC009838 (MH). The authors thank Alexandra Hustedt-Mai for constructive comments on an early version of the manuscript and Kevin Berry for construction of the FPL calibration tubes and custom restraint device.

## Author Contributions

All authors contributed to the study conception and design. Data collection was performed by Samantha Hauser and Andrew Sivaprakasam. Data analysis was done by Samantha Hauser based on analysis code from Hari Bharadwaj. The first draft of the manuscript was written by Samantha Hauser and all authors commented on previous versions of the manuscript. All authors read and approved of the final manuscript.

## Data and Code Availability

Processed data and analysis code are publicly available at https://github.com/hausersn1/sedatedOAEs

## Ethics Approval

All procedures were approved by the Purdue University Institutional Animal Care and Use Committee (Protocol #1110000123)

