## Supplemental Fig 1 for "SEDATION DIFFERENTIALLY AFFECTS DISTORTION-PRODUCT AND STIMULUS-FREQUENCY OTOACOUSTIC EMISSIONS IN CHINCHILLAS"

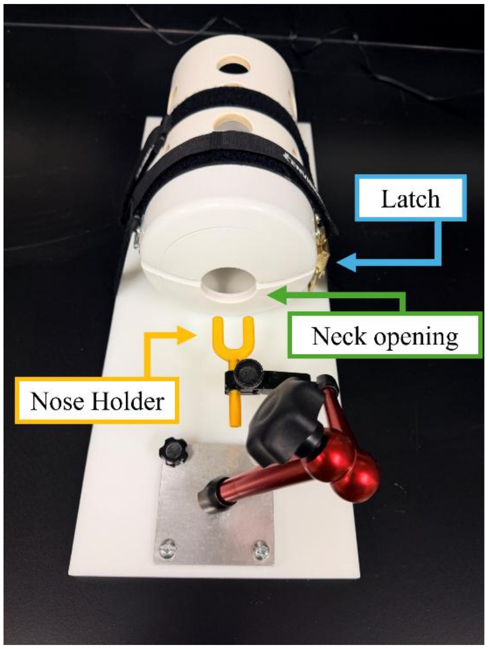


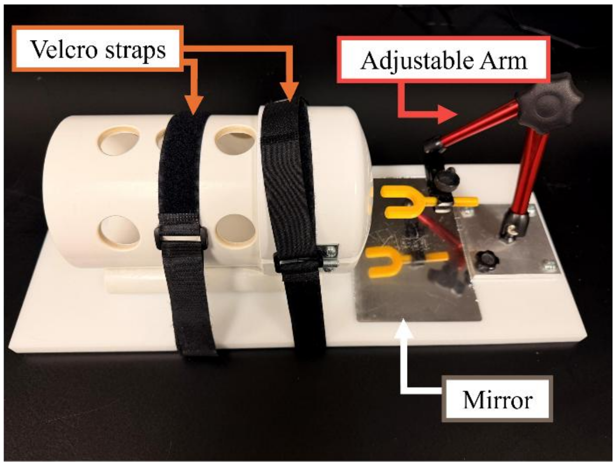


**Online Resource 1** Our chinchilla restraint device for awake measurements improves upon the original design of [29] by including a separate, adjustable nose piece. First, the top half of the cap swings open to allow the chinchilla to enter fully into the tube. The cap is then closed at the neck of the chinchilla and latched. Second, a 3D printed Y-shaped nose holder is adjusted to the correct height and advanced into position at the nasal bridge and secured. This prevents lateral movement of the head while keeping the ears free and the animal comfortable. Velcro straps are used to secure the restraint tube to a plastic board to which the adjustable arm holding the nose piece is mounted. Once positioned, the animals generally remain calm for the duration of testing. A mirror can be placed below the animal's head to monitor breathing and to ensure appropriate positioning.
