## Supplemental Fig 2 for "SEDATION DIFFERENTIALLY AFFECTS DISTORTION-PRODUCT AND STIMULUS-FREQUENCY OTOACOUSTIC EMISSIONS IN CHINCHILLAS"

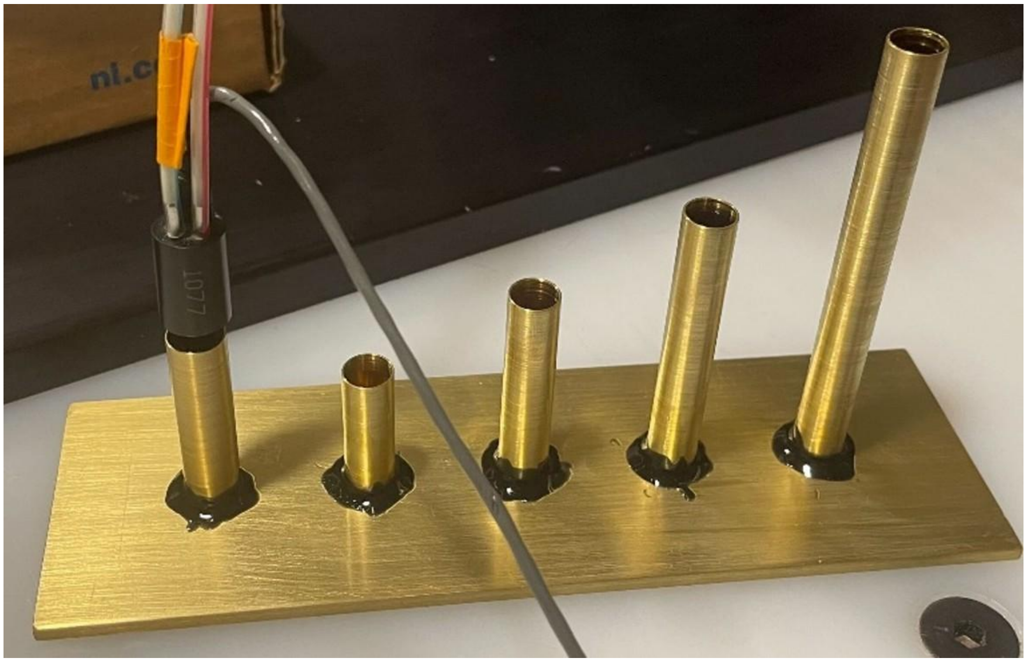

**Online Resource 2** Custom calibration tubes were built with 8 mm inner diameter (i.d.) brass tubing and a brass plate following instructions from [30]. The ER-10B+ probe is shown here positioned in the coupler, which contained 20 mm of the 8 mm i.d. tubing inside of 30 mm of a slightly wider tube (3/8 in outer diameter, shown on the shortest tube). This allowed for quick positioning on each of the five tubes and a single insertion of the probe into the coupler. The tubes had lengths of 18.5, 25.6, 40, 54.3, and 83 mm from left to right.
