## Supplemental Fig 3 for "SEDATION DIFFERENTIALLY AFFECTS DISTORTION-PRODUCT AND STIMULUS-FREQUENCY OTOACOUSTIC EMISSIONS IN CHINCHILLAS"

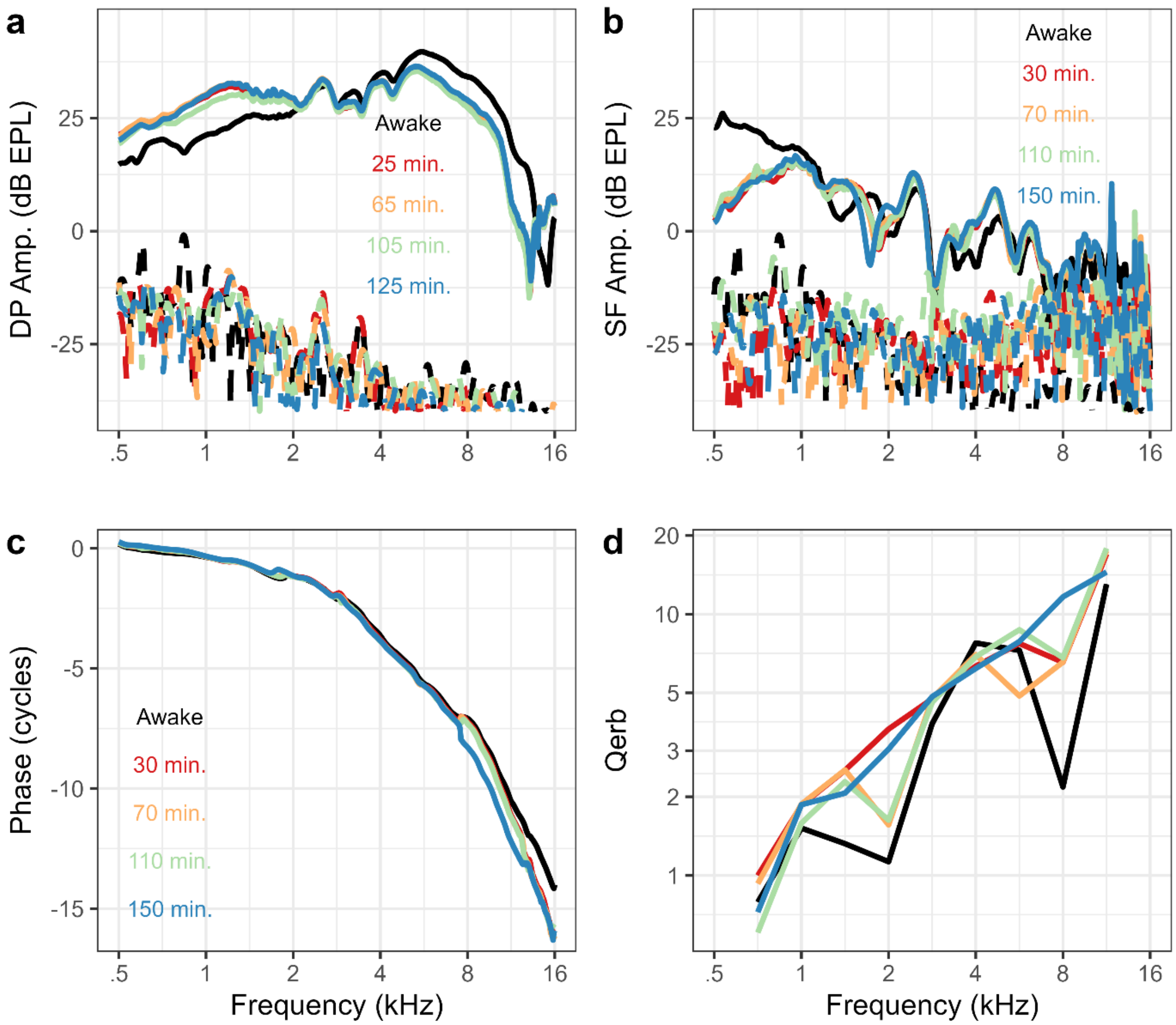


**Online Resource 3** Consistent effects of sedation were observed throughout duration of sedation in one animal. Neither the eartip nor the animal were repositioned during the sedated testing procedure, though it was a different eartip placement from the awake measurement. (a) DPOAE amplitudes are higher when the animal was sedated from .5 to 2 kHz and lower above approximately 3 kHz. (b) SFOAE amplitudes are lower when the animal was sedated below 1 kHz, and higher around 3-4 kHz. SFOAE phase (c) and $Q_{erb}$ estimates (d) show some variation in cochlear tuning estimates across the duration of the experiment, but tuning appears sharper in nearly all sedated measures compared to awake.
